# Sex and area differences in the association between adiposity and lipid profile in Malawi

**DOI:** 10.1101/553776

**Authors:** Ana Luiza G Soares, Louis Banda, Alemayehu Amberbir, Shabbar Jaffar, Crispin Musicha, Alison Price, Moffat J Nyrienda, Deborah A Lawlor, Amelia C Crampin

## Abstract

**Background:** Evidence from high-income countries shows that higher adiposity results in an adverse lipid profile, but it is unclear whether this association is similar in Sub-Saharan African (SSA) populations. This study aimed to assess the association between total and central adiposity measures and lipid profile in Malawi, exploring differences by sex and area of residence (rural/urban).

**Methods:** In this cross-sectional study, data from 12,096 rural and 12,847 urban Malawian residents were used. The associations of body mass index (BMI) and waist-hip ratio (WHR) with fasting lipids (total cholesterol (TC), low density lipoprotein-cholesterol (LDL-C), high density lipoprotein-cholesterol (HDL-C) and triglycerides (TG)) were assessed by area and sex.

**Results:** After adjusting for potential confounders, higher BMI and WHR were linearly associated with increased TC, LDL-C and TG and reduced HDL-C. BMI was more strongly related to fasting lipids than was WHR. The associations of adiposity with adverse lipid profile were stronger in rural compared with urban residents. For instance, one standard deviation increase in BMI was associated with 0.23 mmol/L (95% CI 0.19, 0.26) increase in TC in rural women and 0.13 mmol/L (95% CI 0.11, 0.15) in urban women. Sex differences in the associations between adiposity and lipids were less evident.

**Conclusions:** The consistent associations observed of higher adiposity with adverse lipid profiles in men and women living in rural and urban areas of Malawi highlight the emerging adverse cardio-metabolic epidemic in this poor population. Our findings underline the potential utility of BMI in estimating cardiovascular risk and highlight the need for greater investment to understand the long-term health outcomes of obesity and adverse lipid profiles and the extent to which lifestyle changes and treatments effectively prevent and modify adverse cardio-metabolic outcomes.

## INTRODUCTION

In sub-Saharan Africa (SSA) the prevalence of overweight/obesity is increasing rapidly^1^ in rural and urban populations that are experiencing lifestyle changes (i.e. greater sedentary behaviour, tobacco smoking, alcohol consumption, unhealthy diets and urbanization)^2–6^ and increasing life expectancy (due to reductions in child mortality^7^ and adult HIV-related mortality).^8^ However, the extent to which this transition is affecting the risk and burden of dyslipidaemia is little understood. Although recent Mendelian randomization (using genetic variants as instrumental variables) and randomised controlled trial (RCT) evidence challenge the role of low levels of high density lipoprotein-cholesterol (HDL-C) in causing coronary heart diseases (CHD), high circulating total cholesterol (TC), low-density lipoprotein-cholesterol (LDL-C) and triglycerides (TG) are key causal risk factors for CHD^9,10^, and are effectively and cost-effectively reduced by statins with clear benefits for primary and secondary CHD prevention. In populations of white European origin, higher body mass index (BMI), waist to hip ratio (WHR) and waist circumference (WC) are associated with higher TC, LDL-C and TG.^11^ In these populations, some studies have reported stronger associations of WHR or WC, than BMI, with CHD.^12,13^ However, in by far the largest study to date which included ~222,000 adults from 58 cohorts in an individual participant meta-analysis, similar magnitudes of association were found for BMI, WC and WHR (hazard ratio for CHD (95% CI) for a 1-standard deviation (SD) greater BMI 1.23 (1.17; 1.29), WC 1.27 (1.20; 1.33) and WHR 1.25 (1.19; 1.31).^14^ In that study, reproducibility was also shown to be greater for BMI than WHR (or WC). Prospective studies of children/adolescents in white European populations also find similar magnitudes of association between WC, WHR, waist-height ratio and BMI with blood pressure, fasting glucose and fasting lipids. ^15,16^ Evidence on the association of adiposity and lipid profile in black SSA populations is limited. Most studies in SSA have been small or focused on HIV-positive individuals, and few have collected fasted blood samples and examined differences by sex and area of residence (urban/rural).^17–20^

Some evidence suggests that African women are more adipose than Europeans,^21^ yet people of African ancestry have also been shown to have a more favourable lipid profile.^22^ In a small cross-sectional study undertaken 20 years ago in Cameroon, higher BMI was associated with higher blood pressure and higher fasting insulin, glucose and TG (other lipids were not assayed) in urban residents, but with lower blood pressure and lower fasting insulin, glucose and TG in rural residents.^17^ The rapid epidemiological transition, and the co-existence of insults such as infection and malnutrition (including undernutrition in early life and emerging overweight and obesity in adulthood),^23^ might influence manifestation of adiposity and its association with lipids in SSA populations that differ from those in European populations and this previous SSA study. For example, in Malawi, a very poor SSA country, affected by childhood undernutrition and food insecurity, nearly 25% of the adult population have raised total cholesterol (≥ 5.0 mmol/l).^24^

A better understanding of the extent to which the adiposity (total and central) associates with lipids in SSA, and whether any associations differ by sex and residency (urban/rural) is essential to inform the long-term impact of obesity and dyslipidaemia on the burden of CHD in SSA countries and the development of tools (including cardiovascular risk scores) and interventions appropriate for the setting.

Our aim was to explore whether there was evidence for a causal effect of BMI and WHR on adverse lipid profiles in a poor SSA country and whether any effect differed between women and men and by rural and urban areas. We acknowledge that the data that we are using (population representative data from urban and rural Malawi) is cross-sectional and likely biased by residual confounding (discussed in the limitations of the study). We have carefully considered and controlled for observed confounders, including several measures of socioeconomic position, and believe residual confounding will largely be due to reporting bias of behaviours, such as smoking and physical activity, and our inability to adjust for diet.

## METHODS

### Study design, setting and participants

The Malawi Epidemiology and Intervention Research Unit (MEIRU) collected baseline non-communicable disease data, including interviewer-led questionnaires, anthropometric and blood pressure measures and a fasted blood sample in urban and rural populations using a cross-sectional study design between May 2013 and April 2017. The rural site was in Southern Karonga district, where an established Health and Demographic Health Surveillance Site (HDSS) is located, and the urban site was an enumerated zone in Area 25 of Lilongwe, the capital city, in the Central Region. The two study sites have age and sex structures comparable to the national rural and urban populations and are similar to the national average in most socioeconomic, lifestyle and health indicators ^25,26^. Details on recruitment and data collection were previously described.^27–29^

For this study, we excluded people without serum lipid measurements and those who reported not fasting before the blood collection. Ethical review for this study was obtained from the Malawi National Health Sciences Research Committee (NHSRC; protocol #1072) and London School of Hygiene and Tropical Medicine (LSHTM) Ethics Committee (protocol #6303).

Recruitment and study information was collected at the participant’s household by interviewers and nurses. Questionnaires were available in English, Chichewa (the main language of the Central Region) and Chitumbuka (the main language of the Northern Region), and the data were collected using Open Data Kit on Android operating system tablets.

### Data collection and definitions

Serum lipids (TC, LDL-C, HDL-C and TG) were obtained from a morning venepuncture sample, after a minimum 8-hour fast. Non-fasted participants were re-visited once after a second request to fast overnight. The samples were processed on the same day (within an average of 2.6 hours), and all fasting lipids were assayed using the enzymatic method (Beckman Coulter Chemistry Analyser, model AU480). The two laboratories (one each in the rural and urban centres) participated in external (Thistle RSA system) and internal quality controls (exchanging samples between sites for repeat testing).^29^ Inter and intra-batch coefficients of variation were less than 3% for all lipid measures.

TC, LDL-C and TG were considered high if they were ≥ 5.2 mmol/L, 3.4 mmol/L and 1.7 mmol/L, respectively, and HDL-C was considered low if it was < 1.0 mmol/L. ^30^

Anthropometric measures using standard protocols were taken twice, and the mean of the two used in all analyses. Participants were asked to remove shoes, outer clothing and headdresses. Electronic scales (accuracy of 100g) were used to measure weight, portable stadiometers (accuracy of 1mm) were used for height, and waist and hip measurements were taken using a non-stretch metallic tape with a narrow blade and a blank lead-in. WC was measured on bare skin in the narrowest part of the abdomen between the ribs and iliac crest, and hip circumference was measured over light clothing at the widest part of the buttock.

### Potential confounders

We considered ethnicity, household assets score, education, marital status, parity, smoking status, alcohol intake, physical activity, dietary patterns and intake, lipid-lowering medication, and HIV/ antiretroviral therapy (ART) to be confounders based on their known or plausible effects on adiposity and lipids. We were able to adjust for all of these, except dietary characteristics, as data on diet have not been collected in this study.

Ethnicity was assessed in seven groups: Chewa, Tumbuka, Ngoni, Yao, Lomwe, Nkhonde and Other. Household assets score was based on the score of the cumulative value (using local costings) of 15 items (paraffin lamp, radio, mobile phone, land phone, table & chair(s), bed & mattress, bicycle, TV, electric/gas cooker, sofa set, oxcart, fridge, motorbike, car, and cow), and categorized into quintiles within each area. Educational level was assessed in 7 categories (illiterate, no formal education; literate, no formal education; standard 1-5 years, illiterate; standard 1-5 years, literate; standard 6-8 years; secondary; and tertiary), which were collapsed into 5 categories: no formal education; standard 1-5 years; standard 6-8 years; secondary; and tertiary. Marital status was assessed as never married, married, widowed or divorced/separated. Parity (i.e. the number of pregnancies a woman has experienced irrespective of the outcome) was categorised as 0, 1, 2, 3 or ≥4. Cigarette smoking was categorized into never smoker, former smoker (stopped more than 6 months ago), and current smoker (smoked in the last 6 months). Alcohol intake in the previous 12 months and its frequency were categorized into: never, less than once a month, 1-3 days per month, 1-4 days per week, and 5 or more days per week. The International Physical Activity Questionnaire (IPAQ) was used to measure reported number of days and frequency of walking, and moderate and vigorous physical activity, during leisure-time, at work and when commuting. This information was used to calculate Metabolic Equivalent of Task (MET) values and from these, physical activity was categorized into low, moderate or high. Use of regular lipid-lowering medication was only asked about in those who reported having a diagnosis of raised cholesterol; it was categorised as: no raised cholesterol diagnosis, raised cholesterol diagnosis not taking medication, and raised cholesterol diagnosis taking medication. HIV status was investigated as well as the use of antiretroviral therapy (ART) in those who reported being HIV positive.

### Statistical Analysis

As age distributions vary by sex and area of residence in this population, age-standardised adiposity, lipid levels and proportions of high (or low HDL-C) lipid levels were compared between women and men in rural and urban areas. Age standardisation was done by direct standardisation using the whole cohort, not separated by sex or area. We examined associations between adiposity and lipids as continuous measures and as binary measures; the binary results were considered additional analyses and are presented in Supplementary material.

We used multiple linear regression to examine the association of both BMI and WHR with each lipid measure, adjusted initially for age and then for potential confounding by ethnicity, household assets, education, marital status, parity, smoking, alcohol, physical activity, lipid-lowering medication, and HIV/ART status. We examined the pattern of association (e.g. linear or non-linear) by inspecting the graphs of mean lipid levels by fifths of BMI and WHR measured in standard deviation (SD) units (by sex and area), and comparing models using fifths of BMI and WHR included as categorical variables to those with them treated as a continuous score using a likelihood ratio test. In the absence of strong evidence for a non-linear association, our main analysis results are differences in mean lipid concentrations (mmol/l) for a 1-SD higher BMI or WHR. We accounted for potential clustering (since the primary sampling unit was the household, and all adults in the household were included) by using robust standard errors in all regression models.

Multiple logistic regression analyses were used to examine the association of both BMI and WHR (SD) with each dyslipidaemia, adjusted initially for age and then for other potential confounders. The same confounders used in the linear regression analyses were used for the logistic regressions.

All associations were undertaken in four strata – women from the rural area, men from the rural area, women from the urban area and men from the urban area, as rural and urban residents have different lifestyles. Evidence for differences by sex or area of residence were determined by comparing results across these strata and computing statistical tests for interactions between sex and each anthropometric measurement and between area of residence and each anthropometric measurement. We further explored whether HIV status modified the association between BMI/WHR and lipids ^31^ by computing tests for interactions between HIV status and each anthropometric measurement.

The analyses were carried out in the software Stata 15.1® (Statcorp, College Station, TX, USA).

## RESULTS

From the 33,177 eligible people aged 18 years and above, 30,575 (13,903 in rural and 16,672 in urban area) provided written informed study consent. Fasting lipid measurements were available for 24,943 individuals (12,096 rural and 12,847 urban setting) (Figure 1). In general, those with missing lipid data were more likely to be men, from the urban area, younger, to have lower education, lower assets score, lower BMI, and to be unmarried (Supplementary Tables 1 and 2). Women with missing data were also more likely to have lower parity and lower levels of physical activity.

**Figure 1.**
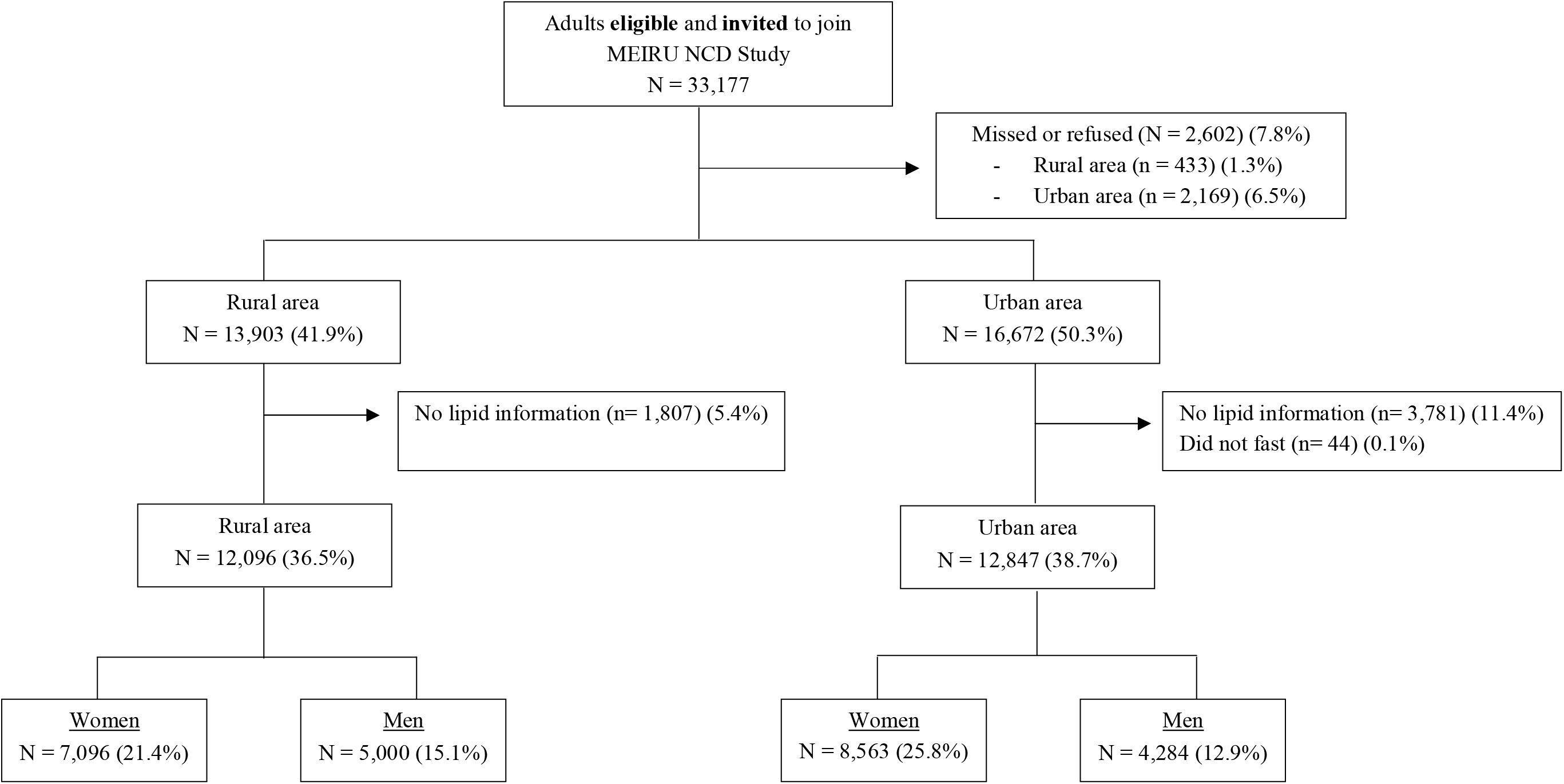
STrengthening the Reporting of OBservational studies in Epidemiology (STROBE) flow chart of the study participants. Proportions are calculated based on the total eligible and invited sample

Compared to the rural population, urban residents were younger, had higher education and median household assets score, and were more likely to be unmarried and to have lower parity (Table 1). Smoking and alcohol drinking were predominantly male behaviours, and current smoking and a higher frequency of alcohol intake was most prevalent among rural men. Most of the population reported high levels of physical activity, especially among women. Fewer than 1% had received a diagnosis of high cholesterol, and of those, 85% reported not taking any lipid-lowering medication. HIV prevalence of those consenting to be tested was 10.6% and it was higher in women and rural residents; 90.8% of participants who were HIV positive reported being on ART. Amongst those included in the analysis, the percentage with missing data for any single variable was between 0 to 6%, except for HIV/ART status, which was missing for 18.5%, with a higher percentage with missing HIV/ART status in the rural area (27.5%) (Supplementary Table 3).

**Table 1.**
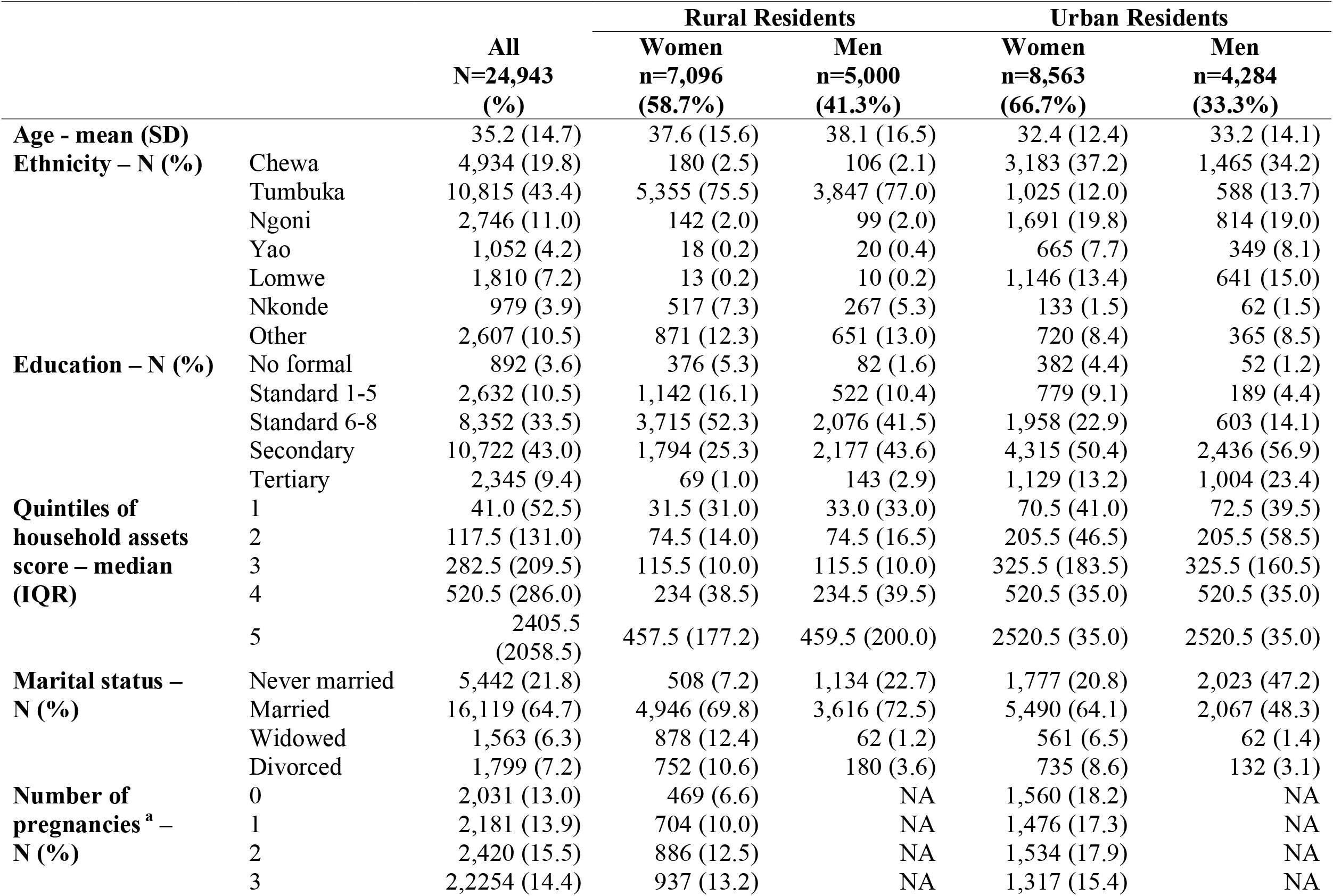
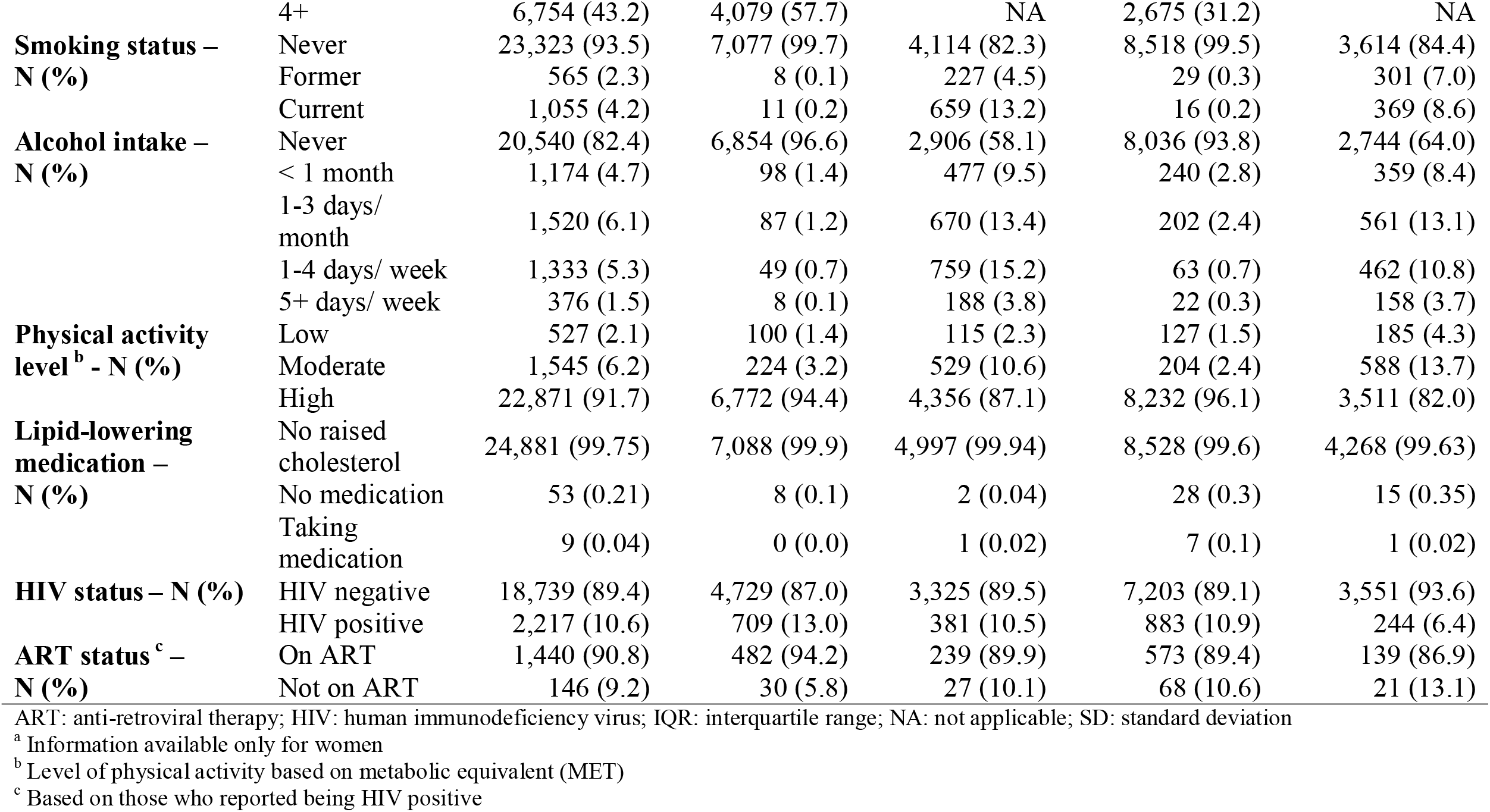
Characteristics of the Malawian population according to area of residence and sex.

BMI was higher in women than men and in urban than rural residents, but WHR was higher in men than women and in rural residents compared to urban residents (Table 2). All age-standardised lipid means were higher in urban than rural participants, with TC, LDL-C and HDL-C being higher in women, and TG higher in men in both rural and urban settings. Of the total participants, 10.8%, 16.4%, 32.9% and 8.9% had high TC, high LDL-C, low HDL-C and high TG, respectively, and 48.9% had any dyslipidaemia, with differences between rural and urban and men and women groups reflecting the pattern of differences in mean levels of these lipids (Supplementary Table 4).

**Table 2.** Age-adjusted mean of body mass index (BMI), waist-hip ratio (WHR) and serum lipids in rural and urban residents, by sex.

|  | **Women**<br>**Mean (95% CI)** | **Men**<br>**Mean (95% CI)** | **p-value** |
| --- | --- | --- | --- |
| **BMI (kg/m<sup>2</sup>)** |  |  |  |
| Rural (N=6,667) / (N=4,990) | 23.3 (23.2; 23.4) | 21.6 (21.5; 21.6) |  |
| Urban (N=8,169) / (N=4,282) | 25.8 (25.7; 25.9) | 22.7 (22.6; 22.8) |  |
| P-value for difference between sex |  |  | <0.001 |
| P-value for difference between areas |  |  | <0.001 |
| **WHR** |  |  |  |
| Rural (N=6,674) / (N=4,990) | 0.85 (0.85; 0.85) | 0.86 (0.86; 0.86) |  |
| Urban (N=8,169) / (N=4,282) | 0.81 (0.81; 0.81) | 0.85 (0.85; 0.85) |  |
| P-value for difference between sex |  |  | <0.001 |
| P-value for difference between areas |  |  | <0.001 |
| **TC (mmol/L)** |  |  |  |
| Rural (N=7,096) / (N=5,000) | 4.00 (3.98; 4.02) | 3.73 (3.70; 3.75) |  |
| Urban (N=8,563) / (N=4,284) | 4.11 (4.08; 4.13) | 3.98 (3.95; 4.01) |  |
| P-value for difference between sex |  |  | <0.001 |
| P-value for difference between areas |  |  | <0.001 |
| **LDL-C (mmol/L)** |  |  |  |
| Rural (N=7,096) / (N=5,000) | 2.62 (2.61; 2.64) | 2.39 (2.37; 2.41) |  |
| Urban (N=8,563) / (N=4,284) | 2.73 (2.71; 2.75) | 2.64 (2.62; 2.67) |  |
| P-value for difference between sex |  |  | <0.001 |
| P-value for difference between areas |  |  | <0.001 |
| **HDL-C (mmol/L)** |  |  |  |
| Rural (N=7,096) / (N=5,000) | 1.15 (1.14; 1.16) | 1.09 (1.08; 1.10) |  |
| Urban (N=8,563) / (N=4,284) | 1.20 (1.19; 1.21) | 1.15 (1.14; 1.16) |  |
| P-value for difference between sex |  |  | <0.001 |
| P-value for difference between areas |  |  | <0.001 |
| **TG (mmol/L)** |  |  |  |
| Rural (N=7,096) / (N=5,000) | 0.85 (0.84; 0.86) | 1.02 (1.00; 1.04) |  |
| Urban (N=8,563) / (N=4,284) | 0.94 (0.93; 0.95) | 1.13 (1.11; 1.16) |  |
| P-value for difference between sex |  |  | <0.001 |
| P-value for difference between areas |  |  | <0.001 |

BMI: body mass index; HDL-C: high density lipoprotein-cholesterol; LDL-C: low density lipoprotein-cholesterol; TC: total cholesterol; TG: triglycerides; WHR: waist-hip ratio

There was no strong evidence of departure from linear associations between BMI/WHR and the lipids in any of the groups (Supplementary Figure 1). Similar patterns of association between the age-adjusted associations (Supplementary Figure 2) and confounder-adjusted analyses were observed. In the confounder-adjusted analyses, there were positive associations of BMI and WHR with TC, LDL-C and TG and inverse with HDL-C in both men and women in rural and urban areas (Table 3, in mmol/L; Figure 2, in SD). The magnitude of the associations, particularly for TC and LDL-C, were stronger for BMI than WHR (e.g., in rural women, 1 SD higher BMI was associated with 0.23 mmol/L (95% CI 0.19, 0.26) higher TC, whilst 1 SD higher WHR was associated with 0.06 mmol/L (95% CI 0.03, 0.09) higher TC). Associations between BMI and serum lipids were mostly similar in magnitude between men and women, but the association between BMI and TG was stronger in urban men than urban women; 1 SD higher BMI was associated with 0.17 mmol/L (95% CI 0.14, 0.20) and 0.09 mmol/L (95% CI 0.08, 0.10) higher TG in urban men and women, respectively. Associations of BMI with TC, LDL-C and TG were stronger in rural than urban women, whereas the associations of BMI with HDL-C and of WHR with TC and LDL-C were stronger in urban than rural women. In men, area differences were less evident in the associations between BMI/WHR and lipids.

**Table 3.** Adjusted association between BMI/WHR and serum lipids in Malawian rural and urban women and men.

|  | **Adjusted difference**<br>**in outcome per 1 SD**<br>**higher BMI (95% CI)** | **Adjusted difference in**<br>**outcome per 1 SD**<br>**higher WHR (95% CI)** |
| --- | --- | --- |
| **Difference in mean TC (mmol/L)** |  |  |
| Rural women (N=4,971) (N=4,975) | 0.23 (0.19; 0.26) | 0.06 (0.03; 0.09) |
| Rural men (N=3,548) (N=3,549) | 0.20 (0.16; 0.23) | 0.07 (0.03; 0.11) |
| Urban women (N=7,455) | 0.13 (0.11; 0.15) | 0.10 (0.07; 0.12) |
| Urban men (N=3,705) | 0.15 (0.12; 0.18) | 0.08 (0.05; 0.11) |
| P-values for difference between sex | 0.012 | 0.442 |
| P-values for difference between area | <0.001 | 0.488 |
| **Difference in mean LDL-C (mmol/L)** |  |  |
| Rural women (N=4,971) (N=4,975) | 0.22 (0.19; 0.24) | 0.06 (0.04; 0.09) |
| Rural men (N=3,548) (N=3,549) | 0.18 (0.16; 0.21) | 0.07 (0.04; 0.11) |
| Urban women (N=7,455) | 0.15 (0.13; 0.17) | 0.10 (0.08; 0.12) |
| Urban men (N=3,705) | 0.17 (0.14; 0.19) | 0.09 (0.07; 0.12) |
| P-values for difference between sex | 0.022 | 0.251 |
| P-values for difference between area | 0.001 | 0.059 |
| **Difference in mean HDL-C (mmol/L)** |  |  |
| Rural women (N=4,971) (N=4,975) | -0.03 (-0.04; -0.02) | -0.03 (-0.04; -0.02) |
| Rural men (N=3,548) (N=3,549) | -0.03 (-0.04; -0.02) | -0.03 (-0.05; -0.02) |
| Urban women (N=7,455) | -0.05 (-0.05; -0.04) | -0.03 (-0.04; -0.02) |
| Urban men (N=3,705) | -0.04 (-0.05; -0.04) | -0.03 (-0.04; -0.03) |
| P-values for difference between sex | 0.603 | 0.730 |
| P-values for difference between area | 0.003 | 0.009 |
| **Difference in mean TG (mmol/L)** |  |  |
| Rural women (N=4,971) (N=4,975) | 0.12 (0.10; 0.13) | 0.11 (0.09; 0.13) |
| Rural men (N=3,548) (N=3,549) | 0.16 (0.13; 0.19) | 0.11 (0.08; 0.14) |
| Urban women (N=7,455) | 0.09 (0.08; 0.10) | 0.09 (0.08; 0.11) |
| Urban men (N=3,705) | 0.17 (0.14; 0.20) | 0.13 (0.11; 0.16) |
| P-values for difference between sex | <0.001 | 0.001 |
| P-values for difference between area | 0.039 | 0.127 |

Serum lipids were assessed in in mmol/L.
BMI: body mass index; HDL-C: high density lipoprotein-cholesterol; LDL-C: low density lipoprotein-cholesterol; TC: total cholesterol; TG: triglycerides; WHR: waist-hip ratio
BMI and WHR are used in standard deviations

**Figure 2.**
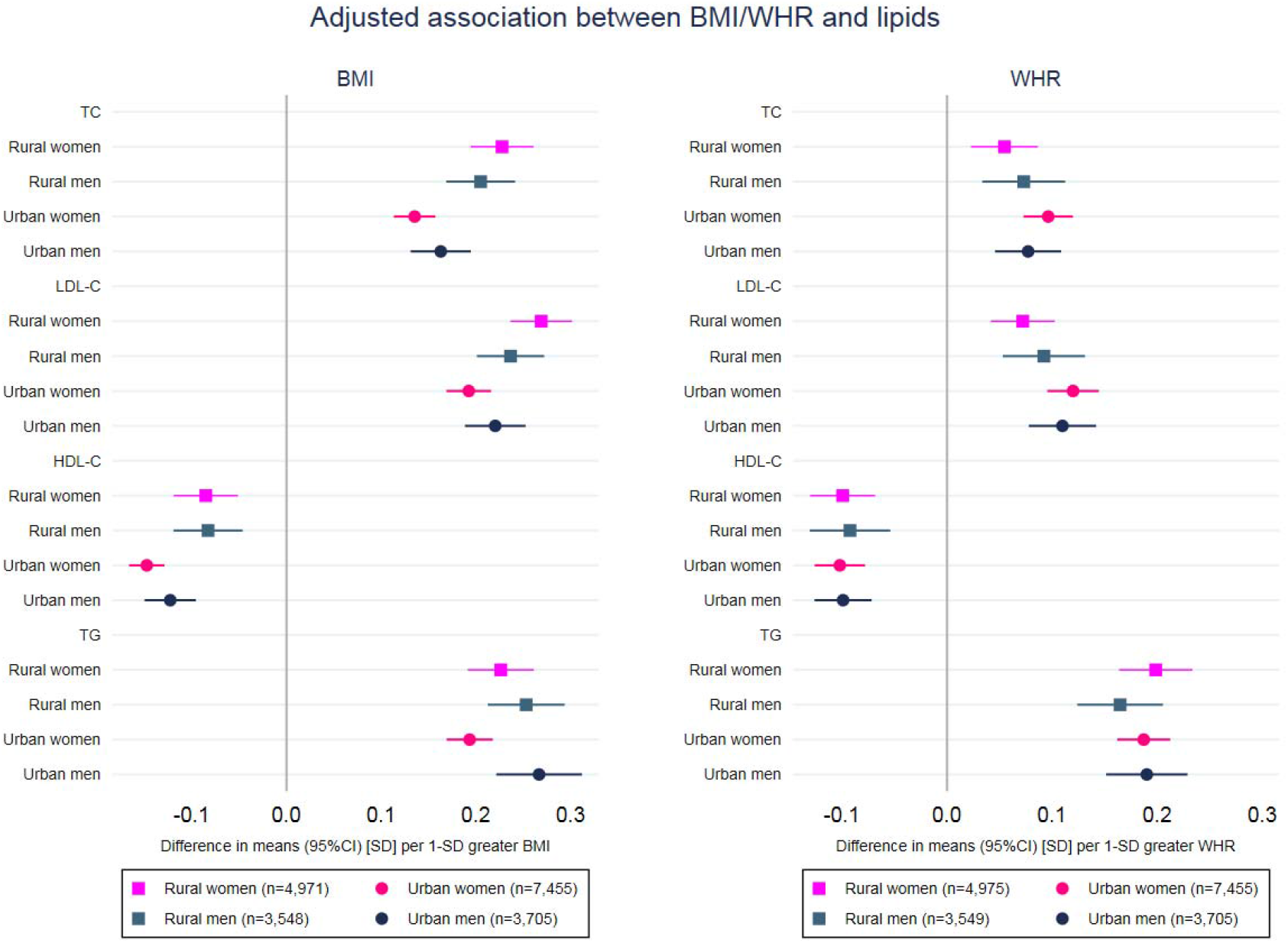
Adjusted association between BMI/WHR and serum lipids in Malawian rural and urban men and women. Adjusted for age, ethnicity, education, household assets score, marital status, parity (women), use of lipid-lowering medication, smoking status, alcohol intake, physical activity, and HIV/ART status. BMI, WHR and serum lipids are used in standard deviations.

Associations with binary levels of each lipid (high TC, high LDL-C, low HDL-C and high TG) followed the patterns we would expect from the results presented here with continuously measured outcomes given associations with BMI/WHR with the continuous measures were linear (Supplementary Table 5, Supplementary Figure 3).

Overall, the associations between BMI/WHR and serum lipids were similar in HIV infected and uninfected individuals (Supplementary Table 6, in mmol/L; Supplementary Figure 4, in SD). Noticeable differences were observed for the association between WHR and TG in women, which was stronger in HIV positive than HIV negative women. For example, in urban women, 1 SD higher WHR was associated with 0.16 mmol/L (95% CI 0.11, 0.22) higher TG in those who were HIV positive and 0.08 mmol/L (95% CI 0.07, 0.10) in those who were HIV negative women.

## DISCUSSION

Our large study in Malawi shows urban residents have a less favourable lipid profile than rural residents and that in both settings women tended to have less favourable profiles than men. After adjusting for several observed confounders, we found positive linear associations of both BMI and WHR with TC, LDL-C and TG, and an inverse association with HDL-C in rural and urban Malawian men and women, with little evidence of differences by sex, but evidence of differences between rural and urban residents. Notably, the association between BMI and all serum lipids, except for HDL-C, was stronger in rural than urban residents, especially in women. The magnitude of the association between adiposity and lipid profile was generally stronger for BMI than WHR. This is consistent with our earlier findings of a stronger associations of BMI than WHR with fasting glucose, diabetes, systolic and diastolic blood pressure, and hypertension,^28^ suggesting that BMI may be a useful predictor of adverse cardio-metabolic outcomes in this population.

Studies have been carried out to identify optimal threshold of anthropometric measures to predict cardiovascular risk in SSA populations and suggested a lower threshold for men and similar or higher threshold for women than those recommended by the World Health Organisation (WHO).^13,32^ Identifying ideal cut-off points of BMI and WHR for dyslipidaemia was out of the scope of this paper. But work to validate cardiovascular risk scores by linking to outcomes in the region is ongoing. Based on the results presented here, we would suggest that prediction models for CVD risk may need to be developed separately for rural and urban SSA populations.

Other studies carried out in SSA countries, including those using WHO STEPS, have also found higher BMI (or higher prevalence of obesity) ^19,33–38^ and a more adverse lipid profile in women than men ^19,20,35,37–39^. Despite differences in adiposity and lipid profiles between men and women, sex differences in the associations between BMI/WHR and lipids were less evident. A markedly stronger association between BMI and TG in urban men than urban women was observed, and further studies to assess whether this replicates in other SSA populations would be useful.

Higher adiposity and poorer lipid profile in urban than rural residents corroborate findings from other studies in SSA. ^34,36,39,40^ Although ethnic differences between rural and urban residents may play a role, in our study results remained unchanged when ethnicity was not included in the confounder-adjusted model (results available from authors on request), and rapid urbanization and its subsequent changes in lifestyle (e.g. decrease in physical activity, and increase in alcohol intake, access to processed food and consumption of unhealthy diets) might explain the more adverse cardiometabolic profile in urban residents compared to those in a rural setting. Even though Malawi has a low percentage of the population living in urban area (about 16%) and has a lower urbanization rate then other SSA countries, the urban population is expected to double by 2050,^6^ and therefore the consequences related to urbanization might become more worrying. A previous study in Uganda has shown that even in rural communities, increasing urbanicity was associated with higher prevalence of lifestyle risk factors for cardiometabolic diseases, such as lower levels of physical activity, higher adiposity and low fruit and vegetable consumption.^41^ Therefore, even small increases in urbanization may have important implications for the burden of adverse lipid profiles in SSA populations.

Stronger associations between BMI and TC, LDL-C and TG, and weaker association with HDL-C were found in rural than urban residents, and this difference was especially observed in women. Ours is by far the largest study of these associations to date. Of previous SSA studies comparing associations of adiposity with lipids between rural and urban communities, one from Cameroon (N = 1,628 participants conducted in 1994) has some consistency with our findings in that there were positive associations of BMI with TG in rural populations, but inverse associations in the urban adults and stronger positive associations between WC and TG in urban than rural participants.^17,42^ A study in Nigerian (N = 247 participants, conducted over 15 years ago) found positive associations of BMI with TC and TG in urban, but not rural, adults.^43^ By contrast, a Ghanaian study (N = 672 participants conducted in 2013) reported stronger associations of WC with TC and LDL-C in rural compared with urban adults.^40^ We have not examined associations of WC with lipids in this paper, but we showed in a previous publication that WC correlated very highly with BMI in the MEIRU cohort.^28^ Thus, this Ghanaian study also has some consistency with our findings. Differences between studies may be explained by differences in the number of participants included, timing in relation to the epidemiological transition and adjustment for confounding factors. However, if the associations of adiposity with lipids are stronger in rural areas as our large study and some other studies suggest, then with urbanization of these areas the CHD epidemic could be particularly marked here.

The prevalence of HIV found in our study is comparable with national estimates, ^26^ and the majority of participants who were HIV positive reported being on ART. HIV infection and ART are known to be associated with cardiometabolic traits, such as adiposity and lipid profile ^31^, and in our study most associations were similar between infected and uninfected participants, but the association of WHR and TG in women was particularly stronger in HIV positive women. We did not find other studies that explored the association between BMI/WHR and lipids in HIV-infected and uninfected individuals is SSA and this single difference by HIV status could be a chance finding.

### Strengths and limitations

Our study was sufficiently large to explore associations by sex and urban and rural locations. The use of fasting lipids and of rigorous, standardised protocols and quality control procedures for data collection are also strengths of our study.

A key limitation of this study is its cross-sectional nature which means we cannot assume that the associations of adiposity with lipids that we have found are causal. Mendelian randomization studies and result of trials of bariatric surgery support causal effects of adiposity on HDL-C and TG, though are less clear for LDL-C.^9,10,44^ We adjusted for a wide-range of plausible confounders, but misclassification of self-reported lifestyle characteristics, such as alcohol consumption and physical activity, might mean that we have not fully accounted for these. We did not have data on dietary intake and therefore could not consider it in the analyses.

The high proportion of missing lipid data in urban residents, especially men, might have resulted in some selection bias and is likely to have underestimated the lipid means and prevalence of dyslipidaemia. The prevalence of high TC, LDL-C and TG was considerably lower than found in a recent meta-analysis of studies in African adults. ^45^ However, our association results would be biased if, in those with missing lipid data, associations were remarkably different from those with lipid information, even after taking into account age, sex, ethnicity, area of residence, education, household assets, smoking status, alcohol intake, physical activity level, and HIV/ART status. Although we cannot rule out this possibility, it is unlikely to be the case. The much higher level of missing data for HIV/ART status in rural compared with urban areas might mean that we have greater residual confounding by this in rural than urban participants. However, this would tend to result in consistently weaker associations in rural groups, which is not what we have found.

### Conclusions and implications

In a large, well-conducted study we have shown that higher BMI and WHR are associated with adverse lipid profiles in men and women in both rural and urban Malawi. Associations were stronger for BMI than WHR, and BMI may be a more reliable indicator of dyslipidaemia than WHR, with greater utility in non-blood based cardiovascular risk score development. Our findings illustrate some of the nature of the emerging cardio-metabolic disease epidemic in Malawi and highlight the need for greater investment to understand the longer-term health outcomes of obesity and adverse lipid profiles and the extent to which lifestyle change and treatments effectively prevent and manage adverse cardio-metabolic outcomes in rural and urban populations in Malawi and potentially in other SSA populations. Efforts for preventing increases in adiposity in both women and men from rural and urban areas should be reinforced in SSA and large prospective studies with detailed baseline data, as presented here, are fundamental to inform scientific and clinical understanding and management of individual risk of CHD in this poor population.

## Acknowledgments

We are grateful to the Karonga and Lilongwe communities, participants and traditional authorities, for their engagement in this work. This study would not have been possible without the MEIRU and Malawi Ministry of Health. We thank the Wellcome Trust for supporting our study.

