## Supplementary Material for "Sex and area differences in the association between adiposity and lipid profile in Malawi"

**CONTENTS**

Supplementary Table 1. Characteristics of the participants with missing and complete lipid data in rural area, by sex (*page 2*)

Supplementary Table 2. Characteristics of the participants with missing and complete lipid data in urban area, by sex (*page 4*)

Supplementary Table 3. Missing data according to area of residence and sex (*page 6*)

Supplementary Table 4. Age-adjusted prevalence of dyslipidaemias in rural and urban residents, by sex (*page 7*)

Supplementary Table 5. Adjusted association between BMI/WHR and dyslipidaemias in rural and urban men and women (page 8)

Supplementary Table 6. Adjusted association between BMI/WHR and serum lipids in Malawian rural and urban men and women, according to HIV status (page 9).

Supplementary Figure 1. Age-adjusted association between quintiles of BMI (a) and WHR (b) with serum lipids in men and women, by area (*page 10*)

Supplementary Figure 2. Age-adjusted association between BMI/WHR and serum lipids in rural and urban men and women (*page 11*)

Supplementary Figure 3. Age-adjusted (a) and fully adjusted (b) associations between BMI/WHR and dyslipidemia in rural and urban men and women (*page 12*)

Supplementary Figure 4. Adjusted associations of BMI (a) and WHR (b) with lipids in HIV positive and HIV negative individuals, in rural and urban men and women (*page 13*)

**Supplementary Table 1.** Characteristics of the participants with missing and complete lipid data in rural area, by sex

|  |  | **Men** | |  | **Women** | |  |
| --- | --- | --- | --- | --- | --- | --- | --- |
|  |  | **Missing** **n=864**  **(14.7%)** | **Non-missing**  **n=5,000**  **(85.3%)** | **p-value** | **Missing** **n=943**  **(11.7%)** | **Non-missing**  **n=7,096**  **(88.3%)** | **p-value** |
| **Age -** mean (SD) |  | 36.2 (16.5) | 38.1 (16.5) | 0.002 | 42.3 (20.6) | 37.6 (15.6) | <0.001 |
| **Household assets score -** median (IQR) | | 179.8 (183.0) | 207.2 (185.0) | 0.008 | 194.0 (195.5) | 201.0 (179.0) | 0.497 |
| **BMI** – mean (SD) | | 21.2 (2.6) | 21.6 (2.8) | <0.001 | 22.5 (4.1) | 23.5 (4.3) | <0.001 |
| **WHR** – mean (SD) | | 0.86 (0.06) | 0.86 (0.05) | 0.105 | 0.86 (0.07) | 0.86 (0.07) | 0.110 |
| **Ethnicity**  % (95% CI) | Chewa | 2.4 (1.6; 3.7) | 2.1 (1.8; 2.6) | 0.354 | 1.9 (1.2; 3.0) | 2.5 (2.2; 2.9) | 0.009 |
|  | Tumbuka | 77.7 (74.8; 80.3) | 76.9 (75.8; 78.1) |  | 70.5 (67.5; 73.3) | 75.5 (74.4; 76.5) |  |
|  | Ngoni | 1.0 (0.5; 2.0) | 2.0 (1.6; 2.4) |  | 2.4 (1.6; 3.6) | 2.0 (1.7; 2.4) |  |
|  | Yao | 0.5 (0.2; 1.2) | 0.4 (0.3; 0.6) |  | 0.5 (0.2; 1.3) | 0.3 (0.2; 0.4) |  |
|  | Lomwe | 0.3 (0.1; 1.1) | 0.2 (0.1; 0.4) |  | 0.2 (0.1; 0.8) | 0.2 (0.1; 0.3) |  |
|  | Nkonde | 6.3 (4.8; 8.1) | 5.3 (4.7; 6.0) |  | 8.6 (7.0; 10.6) | 7.3 (6.7; 7.9) |  |
|  | Other | 11.8 (9.8; 14.1) | 13 (12.1; 14.0) |  | 15.8 (13.6; 18.3) | 12.3 (11.5; 13.1) |  |
| **Education** | No formal | 2.0 (1.2; 3.1) | 1.6 (1.3; 2.0) | 0.065 | 12.7 (10.7; 15.0) | 5.3 (4.8; 5.8) | <0.001 |
| % (95% CI) | Standard 1-5 | 13.1 (11.0; 15.5) | 10.4 (9.6; 11.3) |  | 24.1 (21.4; 26.9) | 16.1 (15.3; 17.0) |  |
|  | Standard 6-8 | 41 (37.7; 44.3) | 41.5 (40.2; 42.9) |  | 39.8 (36.7; 42.9) | 52.4 (51.2; 53.5) |  |
|  | Secondary | 40.3 (37.0; 43.6) | 43.5 (42.2; 44.9) |  | 22.5 (19.9; 25.3) | 25.3 (24.3; 26.3) |  |
|  | Tertiary | 3.7 (2.6; 5.2) | 2.9 (2.4; 3.4) |  | 1.0 (0.5; 1.8) | 1.0 (0.8; 1.2) |  |
| **Marital status**  % (95% CI) | Never married | 26.3 (23.5; 29.4) | 22.7 (21.6; 23.9) | 0.015 | 9.7 (7.9; 11.7) | 7.2 (6.6; 7.8) |  |
|  | Married | 68.2 (65.0; 71.2) | 72.4 (71.2; 73.7) |  | 60.6 (57.4; 63.6) | 69.8 (68.7; 70.9) | <0.001 |
|  | Widowed | 0.7 (0.3; 1.5) | 1.2 (1.0; 1.6) |  | 20.6 (18.1; 23.3) | 12.4 (11.6; 13.2) |  |
|  | Divorced | 4.8 (3.5; 6.4) | 3.6 (3.1; 4.2) |  | 9.2 (7.5; 11.3) | 10.6 (9.9; 11.4) |  |
| **Number of**  **pregnancies ^a^**  % (95% CI) | 0 |  |  |  | 10.0 (8.2; 12.1) | 6.6 (6.1; 7.2) | <0.001 |
|  | 1 |  |  |  | 8.8 (7.2; 10.8) | 10.0 (9.3; 10.7) |  |
|  | 2 |  |  |  | 11.6 (9.7; 13.8) | 12.5 (11.8; 13.3) |  |
|  | 3 |  |  |  | 10.9 (9.0; 13.0) | 13.2 (12.5; 14.1) |  |
|  | 4+ |  |  |  | 58.7 (55.5; 61.8) | 57.7 (56.5; 58.8) |  |
| **Smoking status**  % (95% CI) | Never | 80.7 (77.9; 83.2) | 82.3 (81.2; 83.3) | 0.220 | 99.5 (98.7; 99.8) | 99.7 (99.6; 99.8) | 0.379 |
|  | Former | 4.1 (2.9; 5.6) | 4.5 (4.0; 5.2) |  | 0.2 (0.1; 0.8) | 0.1 (0.1; 0.2) |  |
|  | Current | 15.3 (13.0; 17.8) | 13.2 (12.3; 14.1) |  | 0.3 (0.1; 1.0) | 0.2 (0.1; 0.3) |  |
| **Alcohol intake**  % (95% CI) | Never | 57.6 (54.3; 60.9) | 58.1 (56.7; 59.5) | 0.068 | 95 (93.4; 96.2) | 96.6 (96.1; 97) | 0.038 |
|  | < 1 month | 9.4 (7.6; 11.5) | 9.5 (8.8; 10.4) |  | 1.7 (1.0; 2.8) | 1.4 (1.1; 1.7) |  |
|  | 1-3 days/ month | 11.3 (9.4; 13.6) | 13.4 (12.5; 14.4) |  | 1.6 (1.0; 2.6) | 1.2 (1.0; 1.5) |  |
|  | 1-4 days/ week | 16.1 (13.8; 18.7) | 15.2 (14.2; 16.2) |  | 1.6 (1.0; 2.6) | 0.7 (0.5; 0.9) |  |
|  | 5+ days/ week | 5.6 (4.2; 7.3) | 3.8 (3.3; 4.3) |  | 0.1 (0.0; 0.8) | 0.1 (0.1; 0.2) |  |
| **Physical activity level ^b^** % (95% CI) | Low | 3.5 (2.4; 4.9) | 2.3 (1.9; 2.8) | 0.104 | 5.1 (3.9; 6.7) | 1.4 (1.2; 1.7) | <0.001 |
|  | Moderate | 11.1 (9.2; 13.4) | 10.6 (9.8; 11.5) |  | 5.9 (4.6; 7.6) | 3.2 (2.8; 3.6) |  |
|  | High | 85.4 (82.9; 87.6) | 87.1 (86.2; 88) |  | 89.0 (86.8; 90.8) | 95.4 (94.9; 95.9) |  |
| **Lipid-lowering medication**  % (95% CI) | No raised cholesterol |  | 99.9 (99.8; 100) | 0.772 | 99.7 (99; 99.9) | 99.9 (99.8; 99.9) | 0.109 |
|  | No medication |  | 0 (0.0; 0.2) |  | 0.3 (0.1; 1.0) | 0.1 (0.1; 0.2) |  |
|  | Taking medication |  | 0 (0.0; 0.1) |  | - | - |  |
| **HIV status** | HIV negative | 89.6 (86.7; 91.9) | 89.5 (88.5; 90.5) | 0.975 | 88.4 (85.6; 90.7) | 87.0 (86.0; 87.8) | 0.319 |
| % (95% CI) | HIV positive | 10.4 (8.1; 13.3) | 10.5 (9.5; 11.5) |  | 11.6 (9.3; 14.4) | 13.0 (12.2; 14.0) |  |
| **ART status ^c^**  % (95% CI) | On ART | 5.7 (1.3; 21.4) | 10.2 (7.0; 14.4) | 0.403 | 11.1 (4.0; 27.1) | 5.8 (4.1; 8.2) | 0.197 |
|  | Not on ART | 94.3 (78.6; 98.7) | 89.8 (85.6; 93.0) |  | 88.9 (72.9; 96) | 94.2 (91.8; 95.9) |  |

ART: anti-retroviral therapy; HIV: human immunodeficiency virus; IQR: interquartile range; NA: not applicable; SD: standard deviation

^a^ Information available only for women

^b^ Level of physical activity based on metabolic equivalent (MET)

^c^ Based on those who reported being HIV positive

**Supplementary Table 2.** Characteristics of the participants with missing and complete lipid data in urban area, by sex

|  |  | **Men** | |  | **Women** | |  |
| --- | --- | --- | --- | --- | --- | --- | --- |
|  |  | **Missing** **n=1,520**  **(26.2%)** | **Non-missing**  **n=4,284**  **(73.8%)** | **p-value** | **Missing** **n=2,304**  **(21.2%)** | **Non-missing**  **n=8,563**  **(78.8%)** | **p-value** |
| **Age -** mean (SD) |  | 32.0 (13.0) | 33.2 (14.1) | 0.003 | 30.9 (12.6) | 32.5 (12.4) | <0.001 |
| **Household assets score –** median (IQR) | | 668.2 (358.0) | 793.1 (410.0) | <0.001 | 650.0 (370.0) | 707.4 (436.0) | 0.006 |
| **BMI** – mean (SD) | | 22.2 (3.5) | 22.5 (3.6) | 0.002 | 24.7 (5.0) | 25.6 (5.4) | <0.001 |
| **WHR** – mean (SD) | | 0.84 (0.06) | 0.84 (0.07) | 0.447 | 0.81 (0.07) | 0.81 (0.07) | 0.025 |
| **Ethnicity**  % (95% CI) | Chewa | 39.7 (37.3; 42.2) | 34.2 (32.8; 35.6) | 0.002 | 40.2 (38.2; 42.3) | 37.2 (36.2; 38.2) | 0.034 |
|  | Tumbuka | 11.3 (9.8; 12.9) | 13.7 (12.7; 14.8) |  | 12 (10.8; 13.4) | 12 (11.3; 12.7) |  |
|  | Ngoni | 17.7 (15.9; 19.7) | 19 (17.9; 20.2) |  | 18.8 (17.3; 20.5) | 19.7 (18.9; 20.6) |  |
|  | Yao | 7.6 (6.4; 9.1) | 8.1 (7.4; 9.0) |  | 8.0 (7.0; 9.2) | 7.8 (7.2; 8.4) |  |
|  | Lomwe | 15.1 (13.4; 17) | 15 (13.9; 16.1) |  | 11.8 (10.5; 13.2) | 13.4 (12.7; 14.1) |  |
|  | Nkonde | 1.8 (1.2; 2.6) | 1.4 (1.1; 1.9) |  | 1.0 (0.6; 1.4) | 1.6 (1.3; 1.8) |  |
|  | Other | 6.8 (5.7; 8.2) | 8.5 (7.7; 9.4) |  | 8.1 (7.1; 9.3) | 8.4 (7.8; 9.0) |  |
| **Education** | No formal | 1.7 (1.2; 2.5) | 1.2 (0.9; 1.6) | 0.009 | 5.4 (4.5; 6.4) | 4.5 (4.0; 4.9) | <0.001 |
| % (95% CI) | Standard 1-5 | 5.5 (4.4; 6.7) | 4.4 (3.8; 5.1) |  | 12.1 (10.8; 13.5) | 9.1 (8.5; 9.7) |  |
|  | Standard 6-8 | 13.9 (12.3; 15.8) | 14.1 (13.1; 15.2) |  | 23.0 (21.3; 24.7) | 22.9 (22.0; 23.8) |  |
|  | Secondary | 59.3 (56.8; 61.8) | 56.9 (55.4; 58.3) |  | 45.5 (43.5; 47.5) | 50.4 (49.3; 51.5) |  |
|  | Tertiary | 19.5 (17.6; 21.6) | 23.4 (22.2; 24.7) |  | 14.1 (12.7; 15.5) | 13.2 (12.5; 13.9) |  |
| **Marital status**  % (95% CI) | Never married | 45.8 (43.3; 48.3) | 47.2 (45.7; 48.7) | 0.153 | 23.7 (22.0; 25.4) | 20.8 (19.9; 21.6) | 0.018 |
|  | Married | 48.9 (46.4; 51.5) | 48.2 (46.8; 49.8) |  | 61.5 (59.5; 63.4) | 64.1 (63.1; 65.1) |  |
|  | Widowed | 1.1 (0.7; 1.8) | 1.4 (1.1; 1.9) |  | 6.0 (5.1; 7.1) | 6.6 (6.0; 7.1) |  |
|  | Divorced | 4.1 (3.2; 5.3) | 3.1 (2.6; 3.6) |  | 8.9 (7.8; 10.1) | 8.6 (8.0; 9.2) |  |
| **Number of**  **pregnancies ^a^**  % (95% CI) | 0 |  |  |  | 21.0 (19.4; 22.7) | 18.2 (17.4; 19.1) | <0.001 |
|  | 1 |  |  |  | 20.4 (18.8; 22.1) | 17.2 (16.5; 18.1) |  |
|  | 2 |  |  |  | 18.7 (17.2; 20.4) | 17.9 (17.1; 18.7) |  |
|  | 3 |  |  |  | 14.2 (12.8; 15.7) | 15.4 (14.6; 16.2) |  |
|  | 4+ |  |  |  | 25.7 (24.0; 27.6) | 31.2 (30.3; 32.2) |  |
| **Smoking status**  % (95% CI) | Never | 81.7 (79.7; 83.6) | 84.4 (83.2; 85.4) | 0.014 | 98.9 (98.4; 99.3) | 99.5 (99.3; 99.6) | 0.008 |
|  | Former | 9.3 (7.9; 10.8) | 7.0 (6.3; 7.8) |  | 0.6 (0.4; 1.0) | 0.3 (0.2; 0.5) |  |
|  | Current | 9.0 (7.7; 10.6) | 8.6 (7.8; 9.5) |  | 0.5 (0.3; 0.9) | 0.2 (0.1; 0.3) |  |
| **Alcohol intake**  % (95% CI) | Never | 59.1 (56.6; 61.6) | 64.1 (62.6; 65.5) | 0.002 | 93.6 (92.5; 94.5) | 93.8 (93.3; 94.3) | 0.804 |
|  | < 1 month | 8.4 (7.1; 9.9) | 8.4 (7.6; 9.2) |  | 2.6 (2.0; 3.3) | 2.8 (2.5; 3.2) |  |
|  | 1-3 days/ month | 13.8 (12.2; 15.6) | 13.1 (12.1; 14.1) |  | 2.7 (2.1; 3.4) | 2.4 (2.1; 2.7) |  |
|  | 1-4 days/ week | 13.7 (12.0; 15.5) | 10.8 (9.9; 11.7) |  | 0.7 (0.5; 1.2) | 0.7 (0.6; 0.9) |  |
|  | 5+ days/ week | 4.9 (4.0; 6.1) | 3.7 (3.2; 4.3) |  | 0.3 (0.2; 0.7) | 0.3 (0.2; 0.4) |  |
| **Physical activity level ^b^** % (95% CI) | Low | 3.6 (2.8; 4.7) | 4.3 (3.7; 5.0) | 0.162 | 2.2 (1.7; 2.9) | 1.5 (1.2; 1.8) | 0.001 |
|  | Moderate | 12.3 (10.7; 14.1) | 13.7 (12.7; 14.8) |  | 3.5 (2.8; 4.3) | 2.4 (2.1; 2.7) |  |
|  | High | 84.1 (82.2; 85.8) | 82.0 (80.8; 83.1) |  | 94.3 (93.3; 95.2) | 96.1 (95.7; 96.5) |  |
| **Lipid-lowering medication**  % (95% CI) | No raised cholesterol | 99.8 (99.4; 99.9) | 99.6 (99.4; 99.8) | 0.298 | 99.9 (99.6; 100) | 99.6 (99.4; 99.7) | 0.113 |
|  | No medication | 0.1 (0.0; 0.5) | 0.4 (0.2; 0.6) |  | 0.1 (0.0; 0.4) | 0.3 (0.2; 0.5) |  |
|  | Taking medication | 0.1 (0.0; 0.5) | 0.0 (0.0; 0.2) |  | - | 0.1 (0.0; 0.2) |  |
| **HIV status** | HIV negative | 94.9 (93.3; 96.1) | 93.6 (92.7; 94.3) | 0.125 | 92.7 (91.5; 93.8) | 89.1 (88.4; 89.7) | <0.001 |
| % (95% CI) | HIV positive | 5.1 (3.9; 6.7) | 6.4 (5.7; 7.3) |  | 7.3 (6.2; 8.5) | 10.9 (10.3; 11.6) |  |
| **ART status ^c^**  % (95% CI) | On ART | 8.3 (3.0; 20.8) | 13.1 (8.7; 19.4) | 0.371 | 11.3 (6.8; 18.3) | 10.6 (8.4; 13.3) | 0.822 |
|  | Not on ART | 91.7 (79.2; 97.0) | 86.9 (80.6; 91.3) |  | 88.7 (81.7; 93.2) | 89.4 (86.7; 91.6) |  |

ART: anti-retroviral therapy; HIV: human immunodeficiency virus; IQR: interquartile range; NA: not applicable; SD: standard deviation

^a^ Information available only for women

^b^ Level of physical activity based on metabolic equivalent (MET)

^c^ Based on those who reported being HIV positive

**Supplementary Table 3.** Missing data according to area of residence and sex.

|  | **All** **N=24,943** | **Rural Residents** | | **Urban Residents** | |
| --- | --- | --- | --- | --- | --- |
|  |  | **Women** **n=7,096** | **Men** **n=5,000** | **Women** **n=8,563** | **Men** **n=4,284** |
| BMI | 835 (3.3) | 429 (6.0) | 10 (0.2) | 394 (4.6) | 2 (0.05) |
| WHR | 828 (3.3) | 422 (5.9) | 10 (0.2) | 394 (4.6) | 2 (0.05) |
| Marital status – N (%) | 20 (0.1) | 12 (0.2) | 8 (0.2) | 0 | 0 |
| Parity ^a^ – N (%) | 22 (0.1) | 21 (0.3) | NA | 1 (0.01) | NA |
| HIV/ART status – N (%) | 4,618 (18.5) | 1,848 (26.0) | 1,478 (29.6) | 719 (8.4) | 573 (13.4) |

All the other variables had no missing data.

ART: anti-retroviral therapy; BMI: body mass index; HIV: human immunodeficiency virus; NA: not applicable; WHR: waist-hip ratio

^a^ Information considered only for women

**Supplementary Table 4.** Age-adjusted prevalence of dyslipidaemias in rural and urban residents, by sex.

|  | **Women**  **Prevalence**  **(95% CI)** | **Men**  **Prevalence**  **(95% CI)** | **p-values** |
| --- | --- | --- | --- |
| **High TC (≥ 5.2 mmol/L)** |  |  |  |
| Rural (N=7,096) / (N=5,000) | 11.0 (10.3; 11.7) | 6.9 (6.2; 7.5) |  |
| Urban (N=8,563) / (N=4,284) | 13.3 (12.6; 14.1) | 11.3 (10.3; 12.3) |  |
| P-value for difference between sex |  |  | <0.001 |
| P-value for difference between areas |  |  | <0.001 |
| **High LDL-C (≥ 3.4 mmol/L)** |  |  |  |
| Rural (N=7,096) / (N=5,000) | 16.3 (15.5; 17.1) | 9.8 (9.0; 10.6) |  |
| Urban (N=8,563) / (N=4,284) | 20.8 (19.9; 21.7) | 17.7 (16.5; 18.8) |  |
| P-value for difference between sex |  |  | <0.001 |
| P-value for difference between areas |  |  | 0.016 |
| **Low HDL-C (< 1.0 mmol/L)** | | |  |
| Rural (N=7,096) / (N=5,000) | 33.3 (32.2; 34.5) | 42.5 (41.1; 43.9) |  |
| Urban (N=8,563) / (N=4,284) | 25.7 (24.8; 26.7) | 34.4 (32.9; 35.9) |  |
| P-value for difference between sex |  |  | <0.001 |
| P-value for difference between areas |  |  | <0.001 |
| **High TG (≥ 1.7 mmol/L)** |  |  |  |
| Rural (N=7,096) / (N=5,000) | 6.0 (5.5; 6.6) | 10.4 (9.6; 11.3) |  |
| Urban (N=8,563) / (N=4,284) | 8.8 (8.1; 9.4) | 14.1 (13.0; 15.1) |  |
| P-value for difference between sex |  |  | <0.001 |
| P-value for difference between areas |  |  | <0.001 |
| **Any dyslipidaemia** |  |  |  |
| Rural (N=7,096) / (N=5,000) | 48.2 (47.0; 49.4) | 53.6 (52.2; 55.0) |  |
| Urban (N=8,563) / (N=4,284) | 46.3 (45.2; 47.4) | 51.8 (50.3; 53.3) |  |
| P-value for difference between sex |  |  | <0.001 |
| P-value for difference between areas |  |  | <0.001 |

BMI: body mass index; HDL-C: high density lipoprotein-cholesterol; LDL-C: low density lipoprotein-cholesterol; TC: total cholesterol; TG: triglycerides; WHR: waist-hip ratio

**Supplementary Table 5.** Adjusted association between BMI/WHR and dyslipidaemias in rural and urban women and men.

|  | **Adjusted OR per 1 SD higher BMI (95% CI)** | **Adjusted OR per 1 SD higher WHR (95% CI)** |
| --- | --- | --- |
| **OR of high TC (≥ 5.2 mmol/L)** |  |  |
| Rural women (N=4,971) / (N=4,975) | 1.53 (1.39; 1.69) | 1.18 (1.07; 1.31) |
| Rural men (N=3,548) / (N=3,549) | 1.51 (1.31; 1.73) | 1.40 (1.18; 1.66) |
| Urban women (N=7,455) | 1.30 (1.21; 1.40) | 1.24 (1.14; 1.34) |
| Urban men (N=3,705) | 1.50 (1.35; 1.66) | 1.47 (1.33; 1.62) |
| P-values for difference between sex | <0.001 | <0.001 |
| P-values for difference between area | 0.072 | 0.408 |
| **OR of high LDL-C (≥ 3.4 mmol/L)** |  |  |
| Rural women (N=4,971) / (N=4,975) | 1.53 (1.39; 1.69) | 1.20 (1.10; 1.31) |
| Rural men (N=3,548) / (N=3,549) | 1.51 (1.31; 1.73) | 1.42 (1.22; 1.64) |
| Urban women (N=7,455) | 1.30 (1.21; 1.40) | 1.32 (1.23; 1.41) |
| Urban men (N=3,705) | 1.50 (1.35; 1.66) | 1.45 (1.33; 1.58) |
| P-values for difference between sex | 0.001 | <0.001 |
| P-values for difference between area | 0.038 | 0.118 |
| **OR of low HDL-C (< 1.0 mmol/)** | | |
| Rural women (N=4,971) / (N=4,975) | 1.15 (1.07; 1.23) | 1.20 (1.12; 1.28) |
| Rural men (N=3,548) / (N=3,549) | 1.14 (1.05; 1.24) | 1.18 (1.09; 1.29) |
| Urban women (N=7,455) | 1.30 (1.24; 1.37) | 1.27 (1.20; 1.35) |
| Urban men (N=3,705) | 1.24 (1.16; 1.32) | 1.23 (1.14; 1.33) |
| P-values for difference between sex | 0.223 | 0.284 |
| P-values for difference between area | 0.008 | <0.001 |
| **OR of high TG (≥ 1.7 mmol/L)** |  |  |
| Rural women (N=4,971) / (N=4,975) | 1.89 (1.65; 2.16) | 2.05 (1.69; 2.47) |
| Rural men (N=3,548) / (N=3,549) | 2.02 (1.78; 2.28) | 1.67 (1.45; 1.92) |
| Urban women (N=7,455) | 1.60 (1.45; 1.75) | 1.80 (1.63; 1.99) |
| Urban men (N=3,705) | 1.82 (1.64; 2.03) | 1.69 (1.53; 1.88) |
| P-values for difference between sex | <0.001 | 0.720 |
| P-values for difference between area | 0.001 | 0.651 |

BMI: body mass index; HDL-C: high density lipoprotein-cholesterol; LDL-C: low density lipoprotein-cholesterol; SD: standard deviation; OR: odds ratio; TC: total cholesterol; TG: triglycerides; WHR: waist-hip ratio

Adjusted for age, ethnicity, education, household assets score, marital status, use of lipid-lowering medication, smoking status, alcohol intake, physical activity, and HIV/ART status.

**Supplementary Table 6.** Adjusted association between BMI/WHR and serum lipids in Malawian rural and urban women and men, according to HIV status.

|  | **HIV negative** | **HIV positive** | **p-value** |
| --- | --- | --- | --- |
|  | **Adjusted difference in outcome per**  **1 SD higher BMI (95% CI)** | |  |
| **Difference in mean TC (mmol/L)** |  |  |  |
| Rural women (N=4,400) / (N=679) | 0.22 (0.19; 0.26) | 0.25 (0.14; 0.37) | 0.721 |
| Rural men (N=3,237) / (N=381) | 0.19 (0.16; 0.23) | 0.20 (0.05; 0.34) | 0.973 |
| Urban women (N=6,841) / (N=859) | 0.13 (0.11; 0.15) | 0.17 (0.10; 0.24) | 0.238 |
| Urban men (N=3,550) / (N=243) | 0.16 (0.13; 0.19) | 0.14 (0.00; 0.28) | 0.905 |
| **Difference in mean LDL-C (mmol/L)** |  |  |  |
| Rural women (N=4,400) / (N=679) | 0.22 (0.19; 0.25) | 0.20 (0.11; 0.28) | 0.706 |
| Rural men (N=3,237) / (N=381) | 0.19 (0.16; 0.21) | 0.20 (0.09; 0.30) | 0.993 |
| Urban women (N=6,841) / (N=859) | 0.15 (0.13; 0.17) | 0.15 (0.09; 0.20) | 0.922 |
| Urban men (N=3,550) / (N=243) | 0.17 (0.15; 0.20) | 0.16 (0.04; 0.27) | 0.792 |
| **Difference in mean HDL-C (mmol/L)** |  |  |  |
| Rural women (N=4,400) / (N=679) | -0.03 (-0.04; -0.02) | 0.00 (-0.04; 0.03) | 0.465 |
| Rural men (N=3,237) / (N=381) | -0.04 (-0.05; -0.02) | 0.00 (-0.05; 0.05) | 0.104 |
| Urban women (N=6,841) / (N=859) | -0.05 (-0.06; -0.04) | -0.03 (-0.05; -0.01) | 0.049 |
| Urban men (N=3,550) / (N=243) | -0.05 (-0.06; -0.04) | -0.03 (-0.09; 0.02) | 0.449 |
| **Difference in mean TG (mmol/L)** |  |  |  |
| Rural women (N=4,400) / (N=679) | 0.11 (0.10; 0.13) | 0.13 (0.06; 0.19) | 0.392 |
| Rural men (N=3,237) / (N=381) | 0.16 (0.13; 0.19) | 0.04 (-0.07; 0.15) | 0.095 |
| Urban women (N=6,841) / (N=859) | 0.10 (0.08; 0.11) | 0.08 (0.04; 0.12) | 0.965 |
| Urban men (N=3,550) / (N=243) | 0.17 (0.14; 0.20) | 0.02 (-0.15; 0.18) | 0.167 |
|  | **Adjusted difference in outcome per**  **1 SD higher WHR (95% CI)** | |  |
| **Difference in mean TC (mmol/L)** |  |  |  |
| Rural women (N=4,403) / (N=680) | 0.04 (0.01; 0.08) | 0.15 (0.05; 0.24) | 0.367 |
| Rural men (N=3,238) / (N=381) | 0.06 (0.02; 0.10) | 0.17 (0.03; 0.30) | 0.028 |
| Urban women (N=6,841) / (N=859) | 0.10 (0.07; 0.12) | 0.11 (0.04; 0.18) | 0.530 |
| Urban men (N=3,550) / (N=243) | 0.09 (0.06; 0.12) | 0.05 (-0.04; 0.15) | 0.736 |
| **Difference in mean LDL-C (mmol/L)** |  |  |  |
| Rural women (N=4,403) / (N=680) | 0.05 (0.03; 0.08) | 0.11 (0.04; 0.18) | 0.622 |
| Rural men (N=3,238) / (N=381) | 0.06 (0.03; 0.10) | 0.12 (0.02; 0.22) | 0.101 |
| Urban women (N=6,841) / (N=859) | 0.09 (0.07; 0.12) | 0.10 (0.04; 0.16) | 0.801 |
| Urban men (N=3,550) / (N=243) | 0.10 (0.07; 0.12) | 0.04 (-0.03; 0.11) | 0.468 |
| **Difference in mean HDL-C (mmol/L)** |  |  |  |
| Rural women (N=4,403) / (N=680) | -0.03 (-0.04; -0.02) | 0.00 (-0.03; 0.02) | 0.278 |
| Rural men (N=3,238) / (N=381) | -0.03 (-0.05; -0.02) | 0.01 (-0.05; 0.07) | 0.043 |
| Urban women (N=6,841) / (N=859) | -0.03 (-0.04; -0.02) | -0.02 (-0.05; 0.00) | 0.214 |
| Urban men (N=3,550) / (N=243) | -0.03 (-0.04; -0.02) | -0.03 (-0.09; 0.02) | 0.914 |
| **Difference in mean TG (mmol/L)** |  |  |  |
| Rural women (N=4,403) / (N=680) | 0.09 (0.08; 0.11) | 0.14 (0.07; 0.21) | 0.497 |
| Rural men (N=3,238) / (N=381) | 0.11 (0.08; 0.14) | 0.08 (-0.05; 0.21) | 0.774 |
| Urban women (N=6,841) / (N=859) | 0.08 (0.07; 0.10) | 0.16 (0.11; 0.22) | 0.007 |
| Urban men (N=3,550) / (N=243) | 0.14 (0.11; 0.16) | 0.13 (0.01; 0.26) | 0.885 |

p-value for interaction between HIV/status and anthropometry measures

Serum lipids were assessed in in mmol/L.

BMI: body mass index; HDL-C: high density lipoprotein-cholesterol; LDL-C: low density lipoprotein-cholesterol; TC: total cholesterol; TG: triglycerides; WHR: waist-hip ratio

Adjusted for age, ethnicity, education, household assets score, marital status, parity (women) use of lipid-lowering medication, smoking status, alcohol intake, and physical activity.

BMI and WHR are used in standard deviations

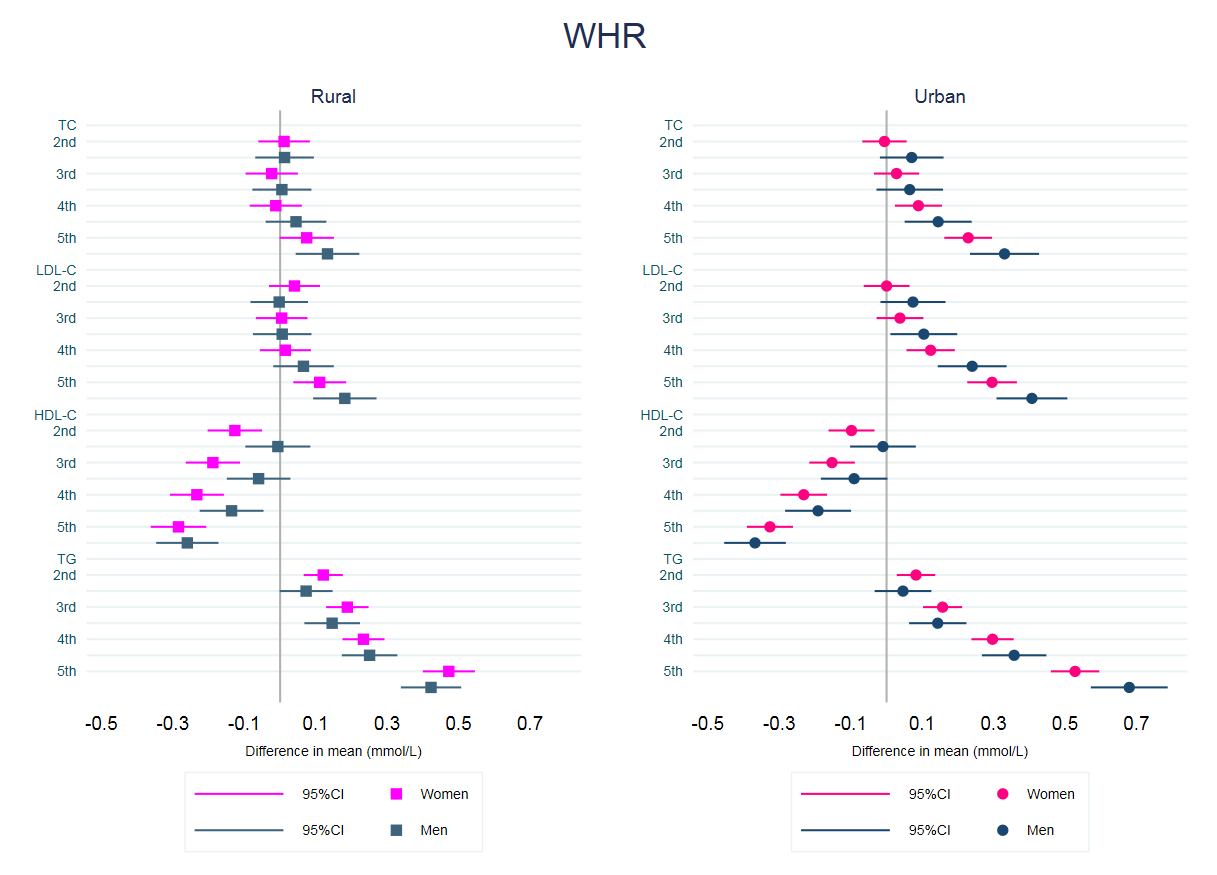

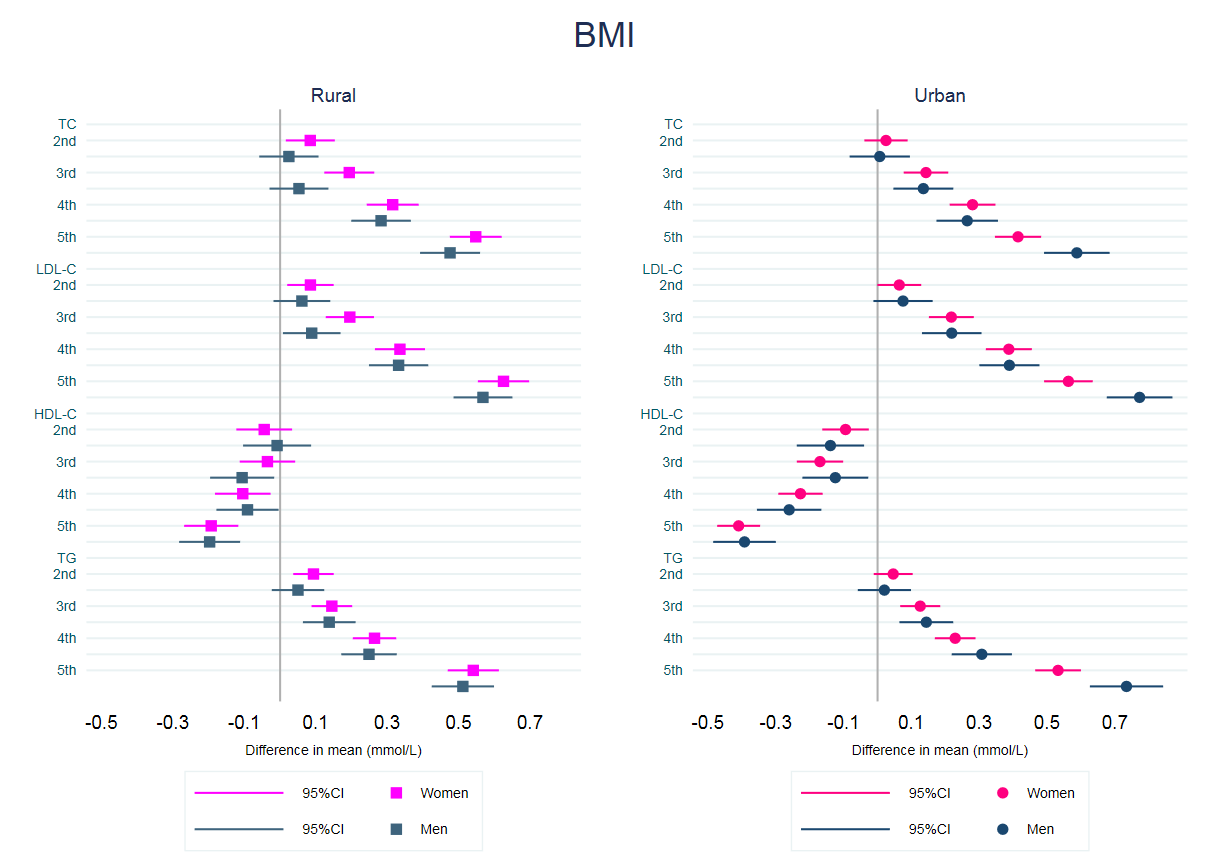

a)

b)

**Supplementary Figure 1.** Age-adjusted association between quintiles of BMI (a) and WHR (b) with serum lipids in men and women, by area.

Serum lipids are used in standard deviations.

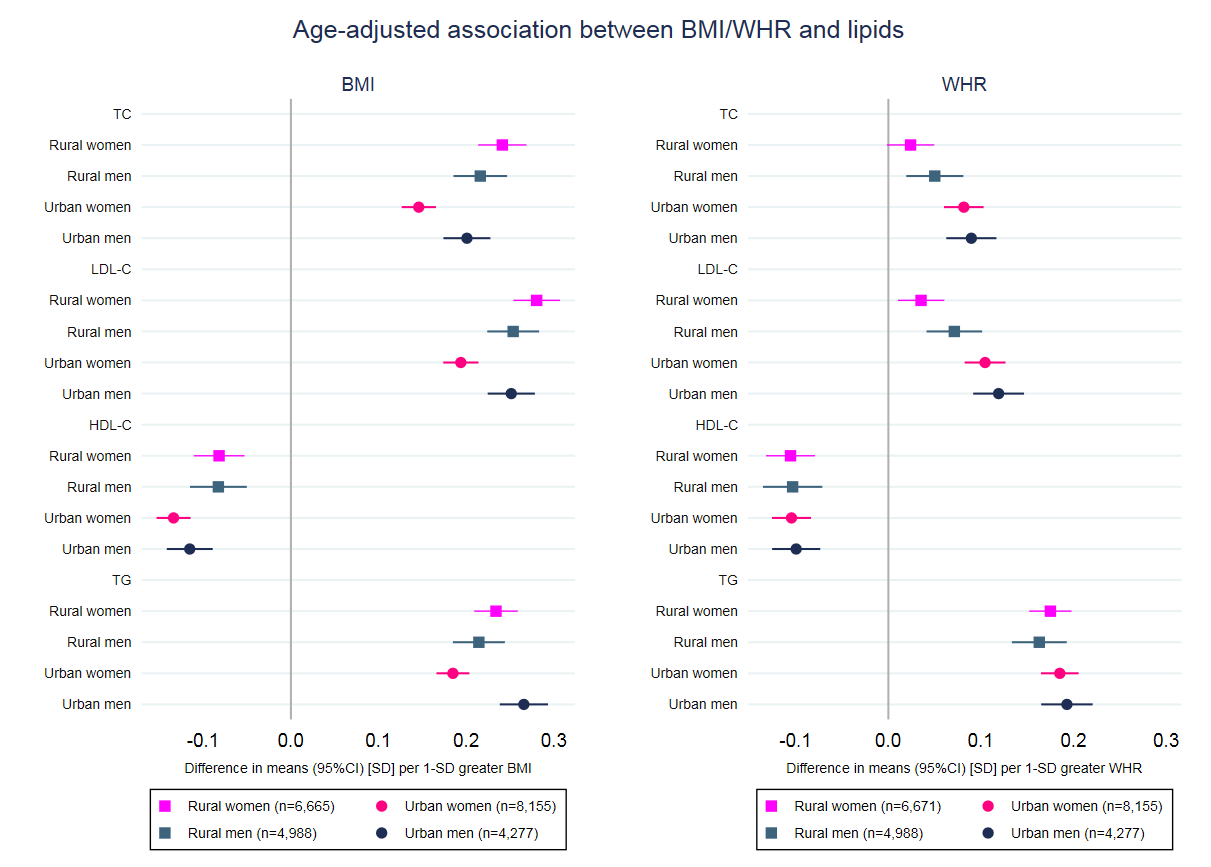

**Supplementary Figure 2.** Age-adjusted association between BMI/WHR and serum lipids in rural and urban men and women.

BMI, WHR and serum lipids are used in standard deviations.

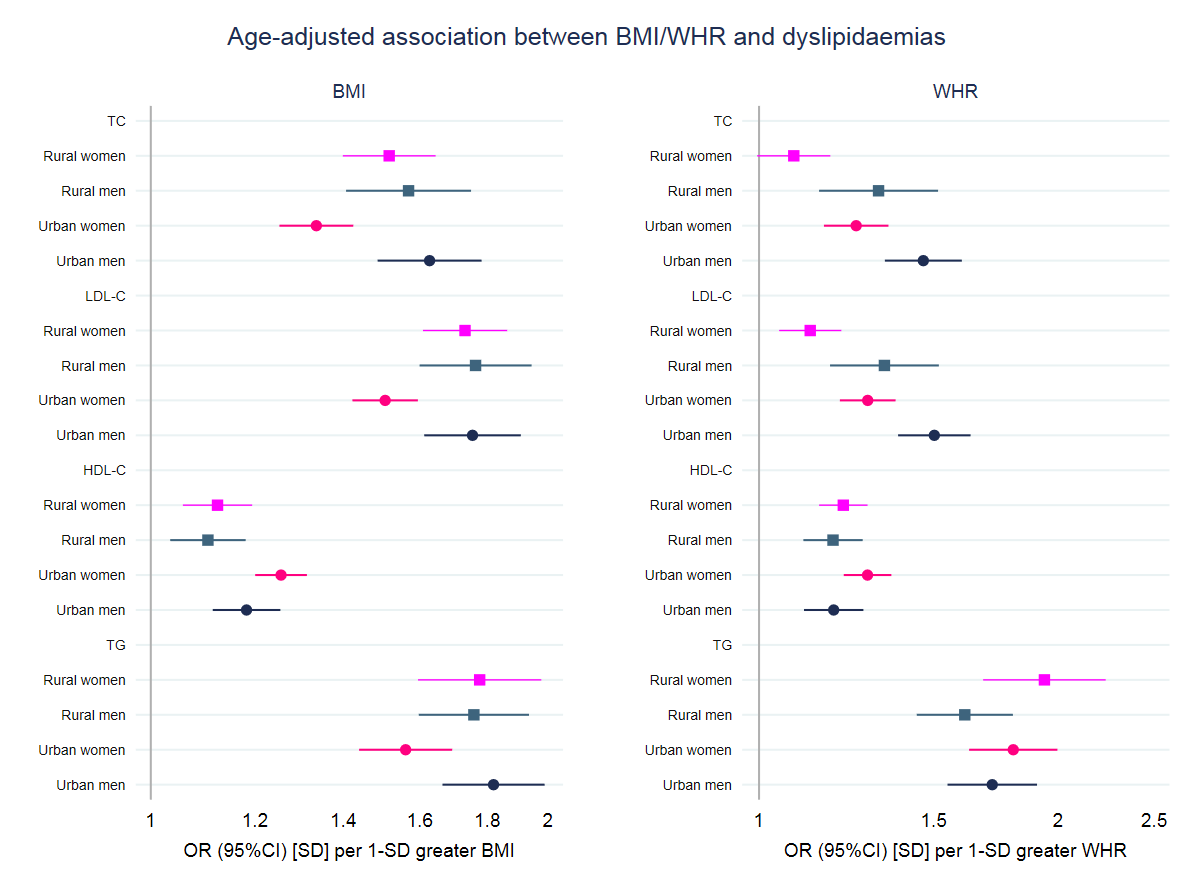

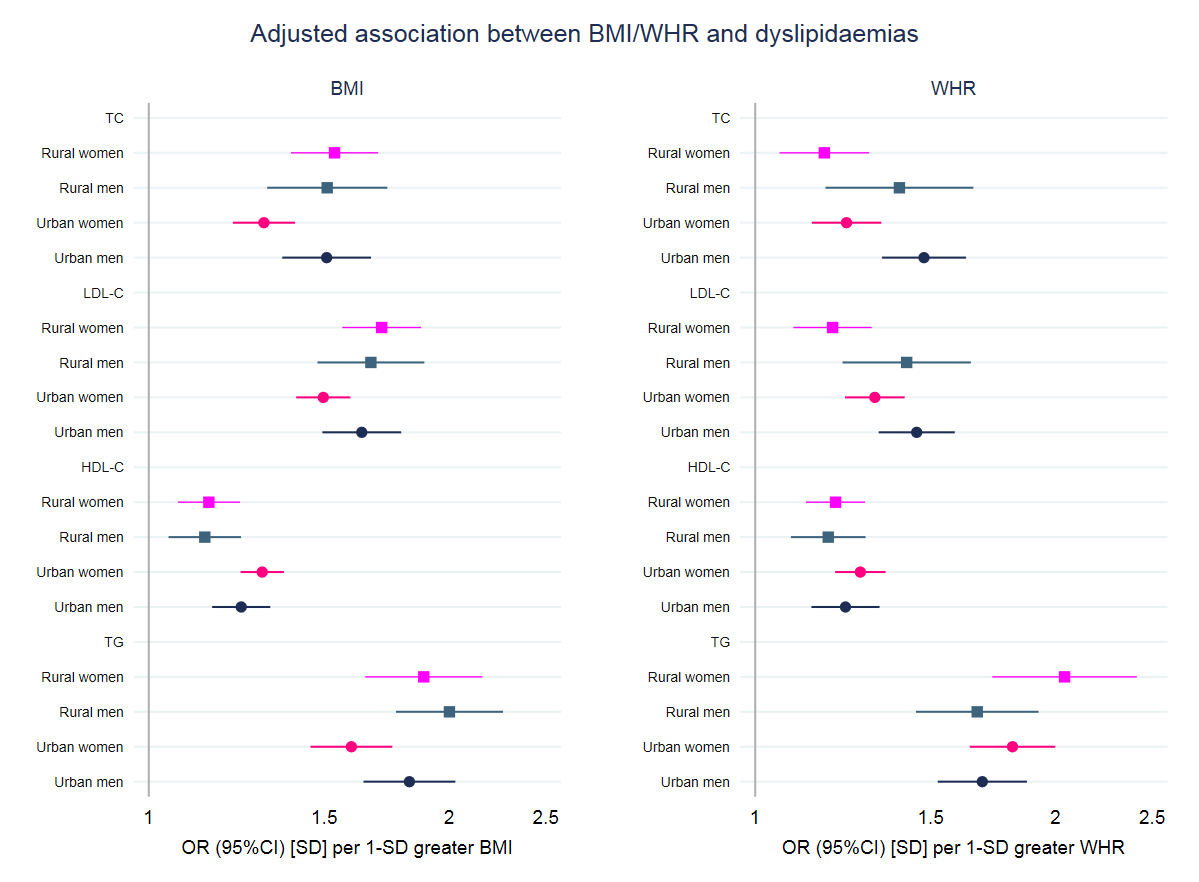

a)

**Supplementary Figure 3.** Age-adjusted (a) and adjusted (b) associations between BMI/WHR and dyslipidaemia in rural and urban men and women.

b)

Adjusted for age, ethnicity, education, household assets score, marital status, use of lipid-lowering medication, smoking status, alcohol intake, physical activity, and HIV/ART status.

BMI, WHR and serum lipids are used in standard deviations.

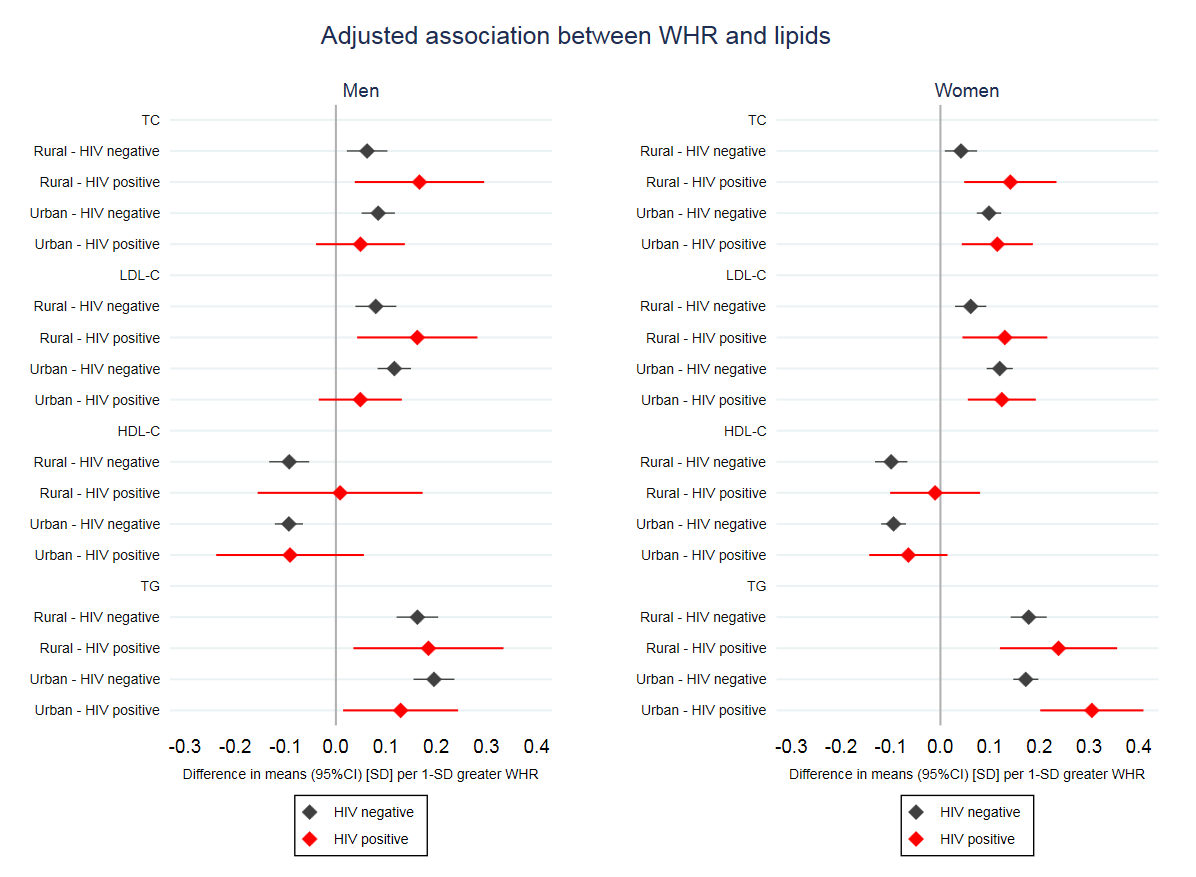

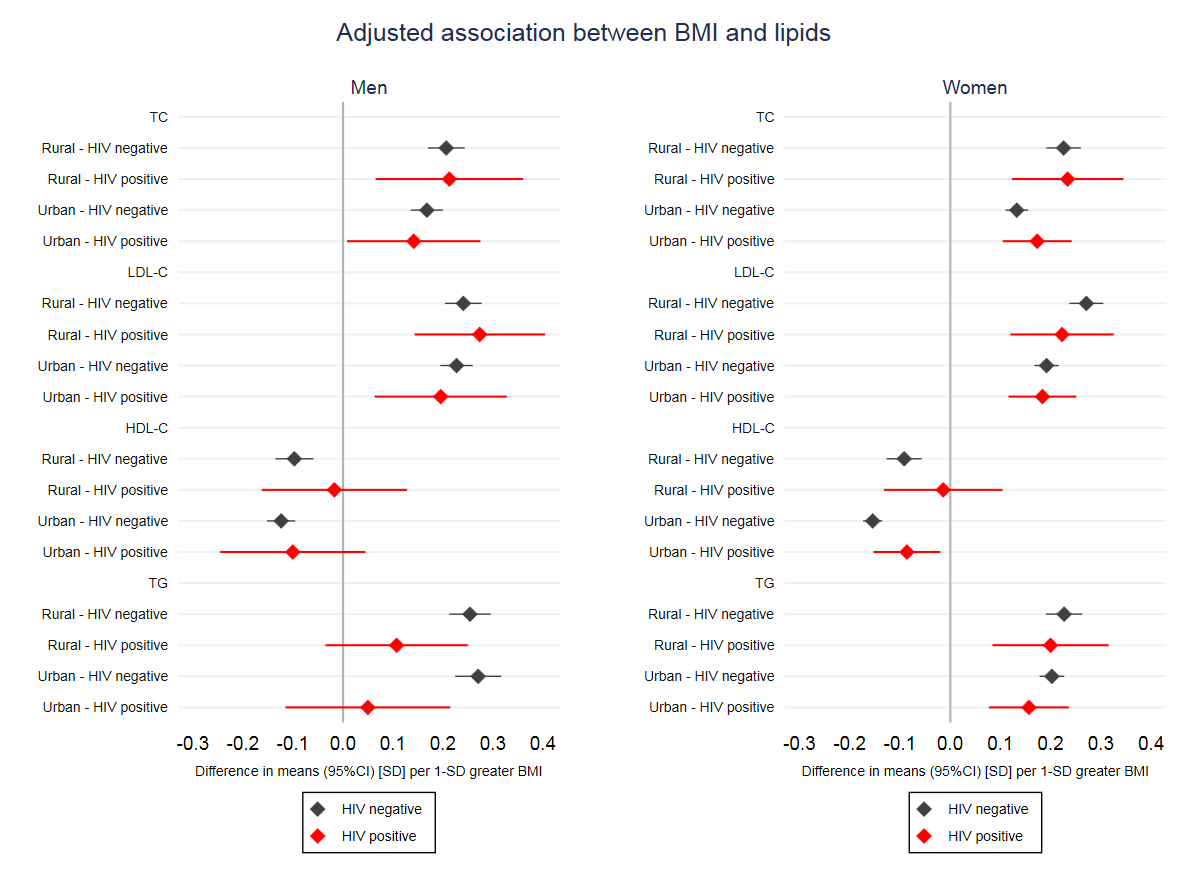

a)

b)

**Supplementary Figure 4.** Adjusted associations of BMI (a) and WHR (b) with lipids in HIV positive and negative individuals in rural and urban men and women.

Adjusted for age, ethnicity, education, household assets score, marital status, use of lipid-lowering medication, smoking status, alcohol intake, and physical activity.

BMI, WHR and serum lipids are used in standard deviations.
